# Temporal genomics reveals rapid parallel adaptation in experimental populations of Trinidadian guppies (*Poecilia reticulata*)

**DOI:** 10.64898/2026.06.15.731539

**Authors:** Ali Hudson, Ronald D. Bassar, David N. Reznick, Joseph Travis, Bonnie A. Fraser

**Affiliations:** Biosciences, Exeter University, Exeter, UK; Department of Biological Sciences, Auburn University, Auburn, AL, USA; Evolution, Ecology, Organismal Biology Department, University of California Riverside, Riverside, CA, USA; Department of Biological Science, Florida State University, Tallahassee, FL, USA

**Keywords:** temporal variation, population genomics, guppy, Poecilia reticulata, rapid adaptation

## Abstract

Understanding the genomic basis of early adaptation is a central question in evolutionary biology. Although there is ongoing debate about whether early adaptation is more likely to be driven by polygenic responses or by loci of large effect, few studies of natural populations have been able to address this problem. Furthermore, early adaptation to a novel environment often coincides with founding events, making it difficult to disentangle neutral effects from adaptive genomic changes. Here we take advantage of the unique *in situ* guppy experimental system, in which we established replicate populations by translocating guppies from a high predation locality to four low predation localities. We present whole genome sequencing from the source population and the four experimental populations, sampled after ∼8-10 generations (first period), and again at ∼18-22 generations (second period). We find signatures of inbreeding only in the first period, despite documented population crashes in the second period. We show genome-wide dynamics of selection as well as selective change at single loci and uncover new targets of selection. Overall, we found signatures of selection at all levels; genome-wide, chromosome, and individual windows are more repeatable among replicates in the first period than in the second period.

## Introduction

How populations adapt to new environments is not only a fundamental question in evolutionary biology but one that has implications for the conservation of species in our rapidly changing world. The early phases of adaptation may be the most important, as it will determine whether a population will persist in its new environment. This is especially true when a founding population in a new environment is small. The intensity of genetic selection necessary for adaptation is inversely proportional to the effective population size, which suggests that early adaptation is more likely to be driven by alleles with large effects on fitness. Alleles with large effects will also facilitate larger steps toward a new fitness peak that is distant from the population’s existing mean trait values (Orr, 1998). However, whether large effect alleles are more commonly selected or are easier to detect remains an open point of debate (Bomblies and Peichel, 2022). Genomic sampling across multiple time points within the early stages of adaptation will help answer this question. Here we investigate early adaptation to a new predation environment in the Trinidadian Guppy (*Poecilia reticulata*) using genomic sampling at multiple time points.

Early phase adaptation may be difficult to detect with only a single snap-shot of genomic diversity, especially when founding populations are small. First, initial colonisation to new environments or sudden changes to the environment are often accompanied by bottlenecks. A single genomic sample may be unable to distinguish effects of subsequent genetic drift from those of novel selection. Second, the responses of quantitative traits even to strong selection can be polygenic. Polygenic responses will be difficult to detect because small allele frequency changes will be distributed across multiple loci. Recent advances in temporal genomics methods, however, can detect autocovariance of allele frequency change across time periods and replicates, providing a powerful new approach in detecting polygenic response (Buffalo and Coop, 2019, 2020).

Experimental populations of guppies offer a unique opportunity to investigate these questions. In the Northern Range mountains of Trinidad, guppies experience a gradient of predation risk, from low risk at high elevations to high risk at low elevations. Studies contrasting the extremes of this continuum have revealed that guppy populations have shown repeated, independent adaptation to low predation (LP) environments. Male guppies from LP sites have larger and more colour spots and increased conspicuousness (Endler, 1980) and mature at a later age and bigger size than males from high predation (HP) environments (Reznick, 1982; Reznick and Endler, 1982). Female guppies from LP sites give birth at a later age and larger size than those from HP sites, and they allocate less resources to reproduction and produce fewer, larger offspring than females from HP sites (Reznick, 1982; Reznick and Endler, 1982). A number of other features distinguish LP from HP guppies, including lower resting metabolic rates (Auer *et al*., 2018), altered head and jaw morphology (Matthews, Reznick and Dial, 2024), and differences in diet (Bassar *et al*., 2010).

Investigators have documented the phenotypic evolution of LP guppies from HP ancestors by translocating HP guppies to an upstream locality that was previously-guppy free, which mimics the natural process through which upstream locations are colonized by guppies. In repeated translocations, guppies have evolved the predicted LP life history and colour in as little as five generations (Reznick *et al*., 1997). These translocations have also revealed that the LP life history phenotype is caused by density-dependent selection, not the direct effect of predation (Bassar *et al*., 2013; Reznick *et al*., 2019; Travis *et al*., 2023).

We have previously offered insight into the genetic architecture of adaptation in guppies. We characterized the early adaptation at the genomic level for four experimentally translocated populations, where a region on chr15 at 5mb shows a repeatable signal of a selective sweep in three of the four experimental populations and parallel change in all four (Van Der Zee *et al*., 2022). We also found signatures of selection in this region in at least two naturally colonised HP-LP river pairs (Fraser *et al*., 2015; Whiting *et al*., 2021). More generally, our comparisons of naturally colonised HP-LP population pairs revealed that specific selected regions vary with river drainage and demographic history, and strong candidates for selection can be found within watersheds (Whiting *et al*., 2021). Through quantitative trait loci (QTL) mapping association studies and large breeding studies, we have also shown that life history traits are likely a combination of polygenic and large-effect loci (Whiting *et al*., 2022). Similarly, male colour pattern is also likely to be oligogenic, with a mix of Y-linked and autosomal loci, although the loci involved may be population specific (Paris *et al*., 2022; Van Der Bijl *et al*., 2025).

Here we used a temporal genomic sampling of replicated experimental guppy populations to explore early phase adaptation to a new environment. We use whole genome sequencing from individuals sampled after ∼8-10 generations (2013) (previously explored in (Van Der Zee *et al*., 2022) and again at ∼18-22 generations (2018). These two time periods roughly correspond to the period in which there was an initial response to selection at high population density (time interval 1: source to 2013) and selection during a steady-state distribution of high densities (time interval 2: 2013 – 2018). We ask 1) whether the signals of population founding and known demographic changes leave genomic signals. 2) Is there evidence of polygenic response to selection and does it differ over time? 3) Is there evidence of selective sweeps and do they differ over time?

### In situ translocation experiment

In 2008 and 2009, we established four replicate “low predation” sites by transplanting guppies from a large, diverse population at a high predation site on the Guanapo river (GH) into four previously guppy-free low predation tributaries upstream (more detail (Reznick *et al*., 2019; Travis *et al*., 2023)). Two translocations were performed in each year; 38 males and 38 females were introduced to the Upper Lalaja (UL) and Lower Lalaja (LL) sites in March of 2008 (resulting in a starting density of 0.18 and 0.14 guppies m^2^ and biomass of 0.027 and 0.016 g/m^2^, respectively). Fifty-two males and females were introduced to the Taylor (TA) and 64 males and females were introduced to the Caigual (CA) tributaries, respectively, in March of 2009 (figure 1) (CA and TA, density of 0.65 and 0.74 guppies m^2^ and biomass 0.105 and 0.136 g/m^2^, respectively).

**Figure 1:**
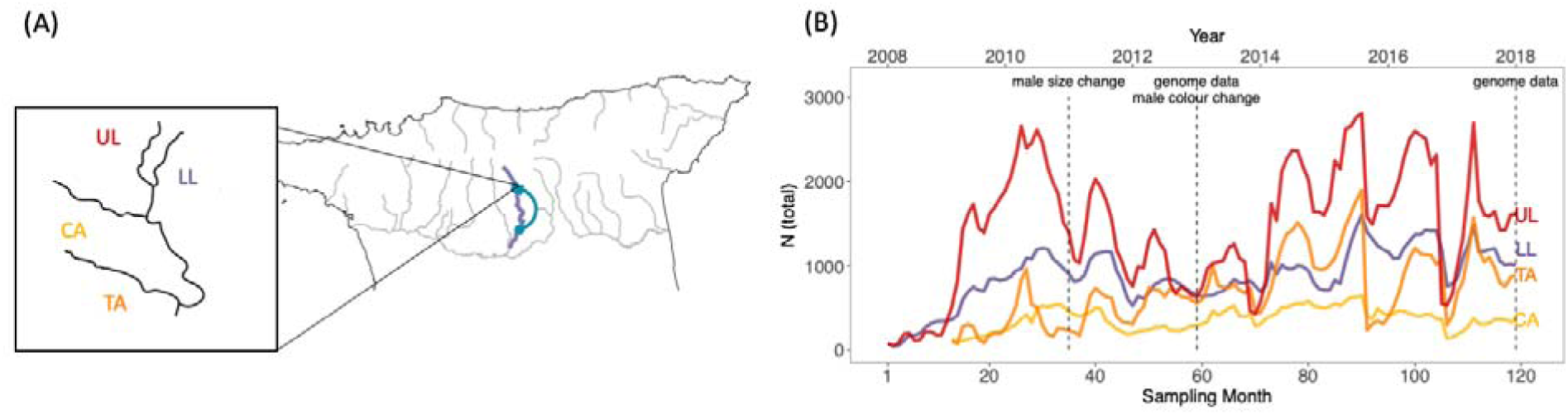
Summary of experimental populations: **A)** Map highlights the Guanapo river in the Northern Range of Trinidad, and an inset shows the four experimental rivers. Arrow indicates the translocation experiment. **B)** Census population size for all populations with dashed lines indicating genomic sampling dates and documented phenotypic changes (Kemp, Batistic and Reznick, 2018; Reznick *et al*., 2019; Travis *et al*., 2023).

We initiated each introduction in a fashion that maximized the genetic variation in each experimental population. Guppies captured from GH as juveniles were reared in single-sex groups, then groups of five males and five females were mated, and the females from each group were introduced to one site and the males to the other. Because females store sperm, all males had the potential to contribute to both introductions within that year, either through mating before translocation and entering as stored sperm or mating after translocation. Natural barriers in the form of waterfalls were reinforced to prevent further migration from downstream guppy populations into the introduction sites. We thinned the forest canopy at the UL and TA sites and left the canopy intact at LL and CA in order to examine the effect of higher productivity in the thinned canopies on guppy dynamics.

We estimated census size and population dynamics with high resolution (appx. 90% probability of capture if alive) monthly mark-recapture censuses (Figure 1) (Travis *et al*., 2023). All four populations displayed rapid initial growth through months 29-32 in the populations established with the smaller number of founders (UL and LL) or months 15-21 for the populations established with the higher number of founders (CA and TA). The populations under thinned canopies (UL and TA) displayed marked oscillations between high densities in the dry seasons and lower densities in the wet seasons. The populations under intact canopies (CA and LL) experienced much milder seasonal oscillations. The populations under thinned canopies (TA and UL) displayed much higher dry-season densities than the populations under intact canopies, although their wet season densities were comparable. All four populations displayed density regulation after the period of initial growth, although TA continued to grow slowly.

All four of the experimental populations have shown predicted heritable phenotypic responses to low predation environments. By 2011 (month 24 for CA and TA, and month 36 for UL and LL) all populations showed an increase in male age and size at maturity compared to the source population in a common garden experiment (Reznick *et al*., 2019). By 2013, Kemp et al. (Kemp, Batistic and Reznick, 2018) found that UL, LL, and TA males had larger coverage of blue/green spots and UL and LL had smaller black spots, compared to the source population.

## Methods

### 2.1 Genomic sampling

In 2013 and 2018, we collected approximately 20 guppies from each experimental site and euthanized and stored them in absolute ethanol at –20°C. The samples are referred to by the population abbreviation and the sample year, e.g. CA13, CA18, while the Guanapo high predation source sample is referred to as GH. GH was also sampled in 2013 and serves as our proxy for starting genetic variation for the experimental populations. DNA was extracted using a QIAGEN DNeasy Blood and Tissue Kit.

We sequenced the GH and 2013 samples to between 7.2x and 11.6x coverage. The GH and majority of the introduction 2013 samples were sequenced with Illumina HiSeq 2000 or 3000 and 250bp paired-end reads, multiplexed with 6-10 fish per lane. An additional 8 samples were sequenced on an Illumina HiSeq 4000 with 150bp paired-end reads. We sequenced all 2018 samples to between 12.7x and 15.2x coverage using an Illumina NovaSeq 6000 to produce 150bp paired-end reads. Sample sizes and coverage details can be found in Table S1.

### 2.2 SNP calling and filtering

We performed SNP sampling following our previous published pipelines (Van Der Zee *et al*., 2022; Whiting *et al*., 2022). We assessed the read quality of 2013 samples with FastQC (Andrews, no date). Low quality bases and adaptors were trimmed with TrimGalore! (Krueger, no date) as part of previous work. We used Fastp (Chen *et al*., 2018) for the 2018 samples to trim sequencing adaptors and remove bases with a phred score <Q22 from the 3’ end of reads and any reads <75bp were discarded. We aligned the trimmed reads to the guppy genome (GCA_904066995.1; (Fraser *et al*., 2020)) with BWA-MEM 0.7.17 (Li and Durbin, 2009). Variants were calibrated with GATK 4.05.1 (McKenna *et al*., 2010) using a truth-set of variants obtained from high-coverage PCR-free sequencing of 12 guppies (Fraser *et al*., 2020).

We filtered the resulting VCFs to retain only biallelic SNPs and remove variants with a QD <2.0, FS >60, MQ <40, a HaplotypeScore >13, MappingQualityRankSum <-12.5, a depth below 5 or above 100. Sites missing from more than 50% of individuals within a population were removed, and finally a minor allele frequency cut-off of 0.01. Individuals with over 35% of missing data were removed from the final VCF.

To enable haplotype-based analyses, we phased SNPs across the whole filtered dataset. Following, Whiting et al. (Whiting *et al*., 2021) we first phased the dataset with Beagle 5.2 (Browning, Zhou and Browning, 2018), then used ExtractPIRs v1 to identify informative reads to pass to SHAPEIT v2(r900) (Delaneau, Zagury and Marchini, 2013), to be phased again. Unless otherwise specified, we used the resulting phased VCF for all downstream analyses. After filtering, 6,350,786 biallelic SNPs remained across a total of 158 individuals (with a minimum of N=10 and maximum of N= 24 per population).

### 2.3 Population structure and diversity

We performed principal component analysis, using Plink 2.0, to examine population structure over time. The PCA was calculated using a linkage pruned dataset; an R^2^ cutoff 0.2, window size of 50 SNPs and a step size of 5 SNPs.

We used the PopGenome R package (Pfeifer *et al*., 2014) to calculate F_ST_, π, and Tajima’s D (TD) in 50kb windows for each population at each sampling time. Similarly, we calculated observed and expected heterozygosity using VCFtools 0.1.15 (Danecek *et al*., 2011) and means were calculated per 50kb window. Whole genome summary stats were calculated before windows were filtered, and Bonferroni-corrected Welch’s t-tests (for Tajima’s D) or Mann-Whitney U tests (for π, heterozygosity) of against the GH source population were conducted. Windows with <50 or >1000 SNPs were excluded from all windowed analyses to keep SNP density comparable.

### 2.4 Runs of Homozygosity

We calculated Runs of Homozygosity (ROH) for each population and year using Plink 1.90 on the phased VCF. The minimum number of SNPs needed to count as an ROH was calculated for each population and year, according to the method developed by Lencz et al. (Lencz *et al*., 2007) and outlined by Purfield et al. (Purfield *et al*., 2017) to minimize false positives while accounting for differences in samples sizes and mean heterozygosity. The minimum length of a run was set to 500kb, and a minimum density of 1 SNP per 50 kb was required for a region to be considered part of a run, with a maximum gap between adjacent SNPs of 1000kb. The scanning window was set to 50 SNPs. A window could contain a maximum of 1 heterozygous SNP and up to 5 missing calls and still be considered part of a run. To identify the ends of a run, the minimum proportion of overlapping windows which needed to be homozygous to call a given SNP part of a run was set to 0.05. Inbreeding coefficients F_ROH_ were calculated as the per-individual sum of ROHs greater than 0.5 divided by the assembly size of 718Mb (the genome size without the sex chromosome, LG12). An MWU test was performed on F_ROH_, comparing each population with the source (GH) and comparing within population between sampling years.

### 2.5 Temporal and Population covariance of genome-wide allele frequencies

Using the filtered, unphased VCF, we calculated the covariance of allele frequency changes between time periods for each replicate population, and the covariance of allele frequency changes between pairs of populations over a given time, using cvtk ver 0.01 (Buffalo and Coop, 2019, 2020). We found no correlation between depth of sequencing and covariance or variance (Figure S1), therefore no bias correction was used. Results were compared to neutral expectations using forward Wright Fisher simulations for each experimental population under neutrality using SLiM5 (Haller, 2026). We modelled evolution on 1 chromosome taken from the GH sample as input and included mutation rate of 4.89e-8 (Künstner *et al*., 2016) and recombination rate of 1e-8 (Whiting *et al*., 2022). We then simulated each population according to the census population data, with a starting population size of 20,000 (N_E_ estimated for GH from (Whiting *et al*., 2021)), and then taking the minimum census size of the population for each generation with standard error sampling. A VCF output was generated for generation 10 and generation 22 (UL and LL) or generation 8, and generation 18 (CA and TA) with sampling the number of individuals used in this study. Simulations were run 100 times to compare to observed data.

### 2.6 Scans for selection at individual loci

Outlier windows were detected if mean allele frequency change was higher than those simulated under neutrality. Simulations were as above and we identified outlier loci as those where allele frequency change surpassed the 95% CI of simulated allele frequency change. Upset plots were used to visualise overlap of outliers using the upset package in R (Conway, Lex and Gehlenborg, 2017).

We then further narrowed our analysis to candidate windows that were outliers in at least 3 of our 4 populations. We identified candidate windows separately for the first interval, the second interval, and the total period. We obtained gene annotation by mapping regions to the ensembl reference genome (GCA_000633615.2) using minimap2. GO enrichment tests were done using the ‘weight01’ algorithm and Fisher’s exact test in TopGO ver (Alexa and Rahnenfuhrer, 2010), with the background set to all genes in the ensembl reference guppy genome. Allele frequency dynamics were examined per population by selecting loci within candidate windows that were above the 95% cut-off simulated allele frequency change per the given population.

## Results

### 3.1 Populations diverge over time and signals of inbreeding are only evident in the first time period

Divergence among the four experimental populations from the GH source population has increased with time. Principal component analysis revealed the 2018 samples to be farther from the source population along the first and second PC axes (Figure 2A). The largest component, PC1 (accounting for 4.47%), separates LL and UL from CA and TA, likely reflecting the different years of initiation and shared starting allele frequency in the members of each pair. PC2 further separates CA samples from TA, and to a lesser extent LL and UL (accounting for 3.24%) (Figure 2A). PC3 and PC4 further separate each population and sampling year but account for a smaller amount of the variation (Figure S2). The other PCs account for little of the variation, each less than 2%.

**Figure 2:**
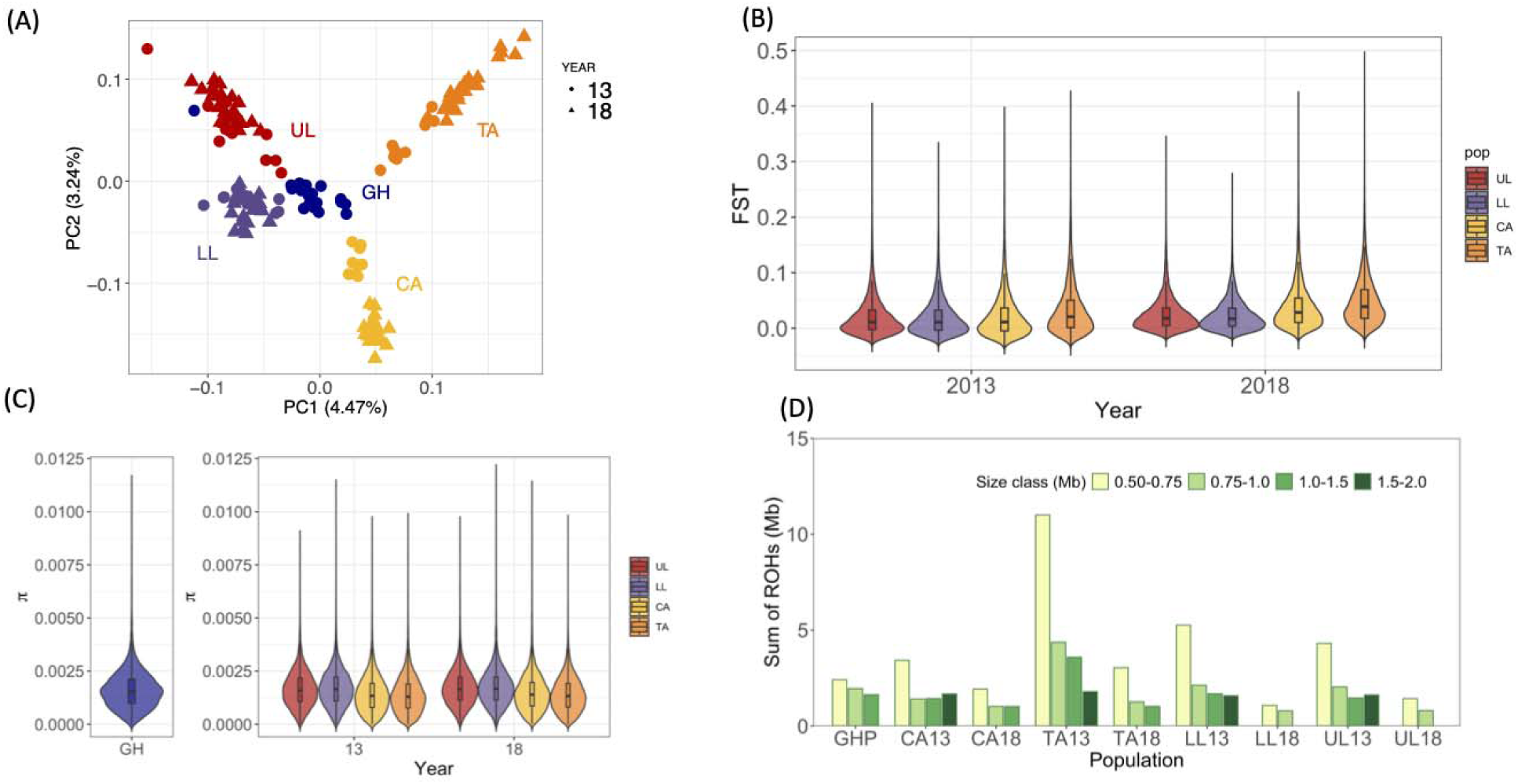
Genome-wide change in experimental populations. **A**. PCA of the source population (GH) and the translocated populations sampled in 2013 and 2018, rivers are indicated by colour and sampling year indicated by shape. **B**. Genome-wide F_ST_ between the source population (GH) and each of the populations per sampling time point. **C** Genome-wide distributions of π (nucleotide diversity) for each population per sampling data. **D**. Sum of ROHs per individual per population distribution of ROHs by size class.

While all four experimental populations continued to diverge from their starting points, CA and TA diverged more after the second sampling than did UL and LL. The mean F_ST_ was small in all comparisons with means ranging between 0.019 (GH – LL13) to 0.049 (GH – TA18) (Figure 2B). Comparing F_ST_ among time intervals, the difference between GH-2013 and GH-2018 was smaller for the UL and LL populations (GH – LL13: 0.019, GH – LL18: 0.024, GH – UL13: 0.020, GH – UL18: 0.025) than the mean F_ST_ for CA and TA, where the mean F_ST_ nearly doubles for CA overtime (GH – CA13: 0.021, GH – CA18: 0.038, GH – TA13: 0.033, GH – TA18: 0.049, Table S2). Across both time intervals, TA has the highest mean F_ST_ when compared to its source, while UL and LL have the lowest. Similarly, comparing populations within time intervals, TA tends to be the largest outlier, with the highest pairwise mean F_ST_ in the first time interval (TA13-UL13: 0.041) and in the second time interval (TA18-CA18:0.066). The two populations with the lowest mean F_ST_ in both time intervals are LL and UL (LL13-UL13: 0.010, LL18-UL18:0.027, Table S3).

The changes in diversity are comparable across all populations and periods (comparing each population’s π, Tajima’s D, expected heterozygosity to the GH source). There was a slight decrease in diversity in CA and TA compared to GH in the first interval but not in the second and UL and LL had similar levels of diversity to the source across both time intervals (Figure 2C: π,: Figure S3:Tajima’s D Figure S4: Expected heterozygosity, Table S4: comparisons with source).

We also tested for changes in levels of ROH (runs of homozygosity); an increase in length and number of ROH is predicted after population bottlenecks. All experimental populations showed the ROHs in the longest size class (1.5 – 2.0 Mb) in 2013, while these were absent in the source GH and in all experimental populations in 2018 (Figure 2D, Figure S5). Comparing F_ROH_ between the source and the 2013 populations, all populations showed a significant increase in F_ROH_ except CA, while we observed no difference between the source and F_ROH_ in 2018, except in LL when the F_ROH_ was significantly smaller than the source (Table S5). Likewise, comparing F_ROH_ between 2013 and 2018 within the population, all populations showed a significant decrease (Table S5).

### 3.2 Polygenic selection was detected during the first time interval via genome-wide allele frequency covariance

During the first time interval, all genome-wide covariances between populations were positive and higher than the mean of simulated neutral. The CA-TA covariance was the highest (observed value was at the 97th percentile of simulated covariances), followed by LL-UL covariances (92%) and TA-UL covariances (92%). All other comparisons were above 85% cut-off (CA-LL: 86%, CA-UL: 88%,TA-LL: 88%) (Figure 3a). Investigating covariance per chromosome revealed that comparisons CA-TA had the highest number of chromosome outliers (CA-TA:12, CA-LL: 2, CA-UL: 4, TA-LL:3, TA-UL: 5, LL-UL:5, Figure S6). Covariance did not correlate with chromosome size for any of the comparisons (Figure S9).

**Figure 3:**
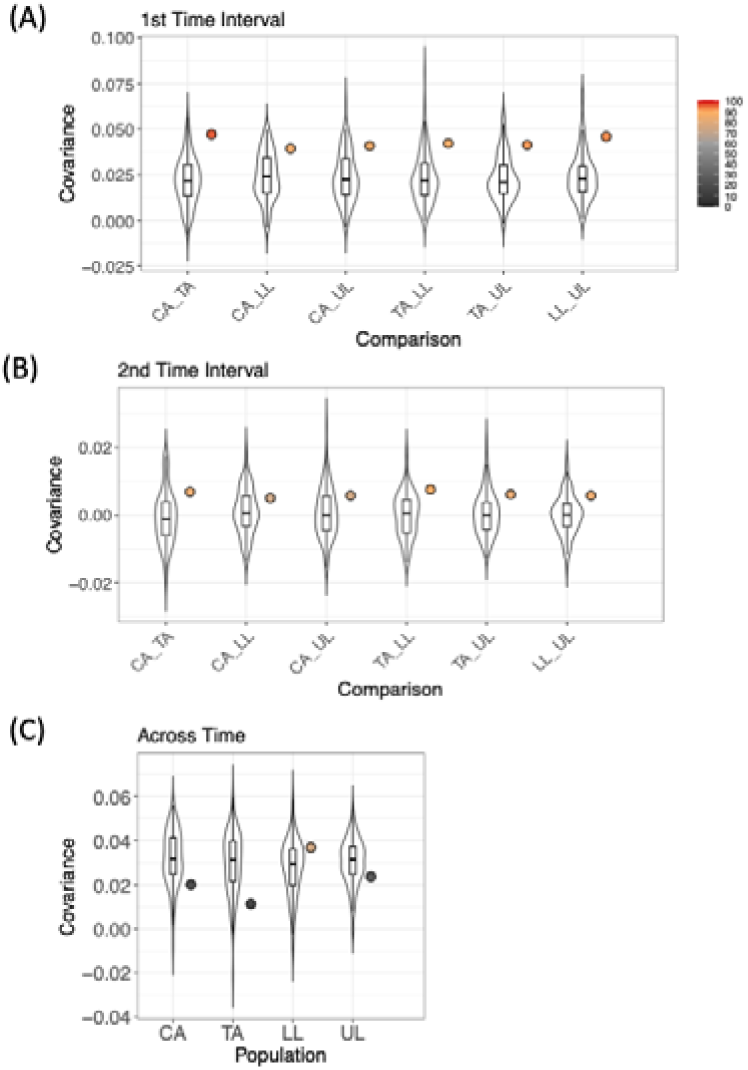
Genome-wide covariance of allele frequency change within and among populations. Violin plots represent the simulated distributions for each comparison under neutral conditions and coloured points are the observed data. Points are coloured by the percentile they fall into within the simulated distribution. **(A)** Covariance between populations within the first time interval (source to 2013) **(B)** Covariance between populations within the second time interval (2013 to 2018) **(C)** Covariances between the two time intervals within one population replicate.

There were chromosomes during the first time interval that showed higher covariance in multiple comparisons. Chromosome 15 was above the 100% of simulated data in 2 comparisons (CA-TA, CA-LL) and above 95% in the remaining comparisons (CA-UL:98%, TA-LL:99%, TA-UL:98%, Figure S6). No other chromosome showed such a significantly repeatable pattern. Indeed, if we remove chr15 from the genome-wide covariance estimates percentiles of observed estimates fall below the 95% percentile in all comparisons but remain high (CA-TA 94%, CA-LL 83%, CA-UL 87%, TA-LL 87%, TA-UL 88%, LL-UL 93%, Figure S10). Only two other chromosomes repeatedly exceed the 95 simulation distribution but not for all comparisons (chr21: CA-TA:99%, CA-LL:95%, CA-UL:89%, TA-LL:98%, TA-UL:100%, LL-UL:96%, and chr1: CA-TA 98%, CA-LL 83%, CA-UL 93%, TA-LL 96%, TA-UL 97%, LL-UL 97%, Figure S6).

During the second time interval, all genome-wide covariances between populations were positive and higher than the mean of simulated neutral covariance but between 73 and 89% quantiles (CA-TA: 84%, CA-LL:73%, CA-UL: 75%, TA-LL: 89%, TA-UL:83%, LL-UL:85% Figure 3b). Covariances per chromosome revealed that chromosome outliers were also lower in all comparisons compared to the first interval (CA-TA:2, CA-LL: 0, CA-UL: 1, TA-LL:6, TA-UL: 4, LL-UL: 4, Figure S7). Accordingly, outlier chromosomes showed less repeatability than in the first time interval, with only chr20 above the 95 percentile in 4 of the 6 comparisons CA-TA: 91%, CA-LL: 54%%, CA-UL:96, TA-LL:100%, TA-UL:98%, LL-UL:99%) (Figure S7).

Finally, covariance between the two time intervals within a given population were below the mean of simulated values for all populations except LL, and here it fell at the 75 percentile (Figure 3c). Similar low covariance was found per chromosome in all populations, with only LL having chromosomes above the 95 percentile (Figure S8). However, chromosomes fell within the lower end of the simulated distribution, below the 0.05 percentile, but these were not consistent among replicates (number of chromosomes below 0.05 percentile: CA: 3, TA: 6, LL: 2, UL: 1).

### 3.3 Signals of selected loci and how their dynamics over time

All four experimental populations showed signals of selected loci in both time intervals and across the whole time period. The majority of observed allele frequency change was within the distribution of simulated neutral allele frequency change, indicating our neutral model captured observed change and outliers are indicative of selection (Figure S11). The percentage of windows that were outliers, defined as falling above the 95% threshold of the simulated distribution per population, were similar across all time intervals but differed among populations. TA consistently had the most outliers (1st time interval: 8.75%, 2nd time interval: 8.48% Total Time period: 8.33%) followed by CA (1st time interval: 5.72%, 2nd time interval: 5.89% Total Time period: 5.01%). While UL and LL had less than half the number of outliers in all time periods (LL: 1st time interval: 2.85%, 2nd time interval: 2.52% Total Time period: 1.45%, UL: 1st time interval: 2.98%, 2nd time interval: 2.74% Total Time period: 1.81% Table S6).

Shared outliers among populations was highest in the first time interval (Figure 4). The number of windows found in all four replicates was 14, with 85 outlier windows shared in at least 3 out of 4 populations. The second time interval showed the least amount of overlap (N=22 in 3 of 4). Finally, 60 windows were shared among the total time interval among 3 of 4 populations.

**Figure 4:**
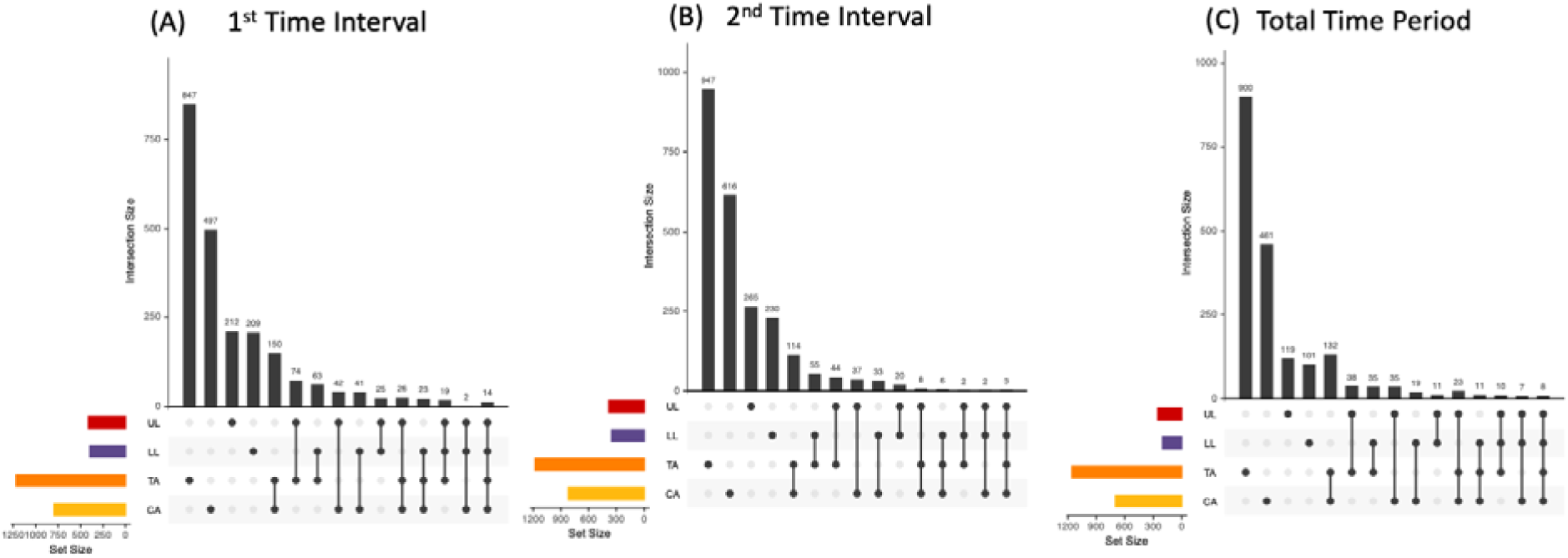
Intersection of outlier windows in allele frequency change for each time interval. (A) Over the first time interval source to 2013 (B) Over the second time interval 2013 to 2018 (C) Over the entire time period source to 2018.

We took windows that were outliers in at least 3 of our 4 populations per time interval as our strongest candidates of selection, and examined them further for gene content and allele frequency dynamics. We refer to these windows hereafter as ‘candidate windows’ (Table S7). These candidate windows were mostly unique to time intervals, where only 1 candidate window was found in time intervals 1 and 2, no candidate windows were shared between time period 2 and total time period, and 17 windows were shared in both time interval 1 and total time period (Table S7).

Examining the gene content of candidate windows revealed possible targets for selection. A full list of gene annotations can be found in Table S8 – Table S10. In the first time interval we found candidate windows that were previously reported and examined for gene content (e.g. chr8:9Mb and chr15:5Mb (Van Der Zee *et al*., 2022)). Interestingly, two windows that were strong candidates in both time interval 1 and total time interval (chr7:18.9Mb and chr7:14.0Mb) contain well known fish pigmentation genes, mitfb (Luo *et al*., 2021) and atp6ap1b (Ramos-Balderas *et al*., 2013). A candidate window in the first time interval (chr4:3.8) contains a well characterised gene involved in the regulation of appetite in humans CARTPT (Yosten *et al*., 2021). Gene Ontology enrichment showed enrichment of biological processes that may be involved in adapting to a new environment (Table S11) including; detection of chemical stimulus involved in sensory perception of smell in time interval 1, regulation of secretion, including cortisol secretion in time interval 2, and retinal pigment epithelium development in total time interval.

We further investigated these candidate windows for allele frequency dynamics (Figure 5, Figures S12-S13, Table S12). Loci within candidate windows identified in the first time interval tended to experience little change in the second, where the distribution of allele frequency change in the second time interval strongly overlapped with zero. This pattern indicates a strong initial sweep and then a plateau or fixation. Conversely, loci within candidate windows identified in the second time interval, tended to experience an opposite but small change in direction as the first, i.e. when allele frequency change was positive in the first time interval it was negative in the second and vice versa. Finally, allele frequency change within candidate windows identified in the total time period tended to be in the same direction in both time intervals, which is expected of a change that is continuous through time. However, in most populations the distribution of allele frequency changes overlaps with zero in the second time interval more than in the first, indicating that the majority of the change happened in the first time interval with moderate or no change happening in the second. Dynamics did not differ substantially between populations most likely because of our filtering of overlapping outliers to identify candidates.

**Figure 5:**
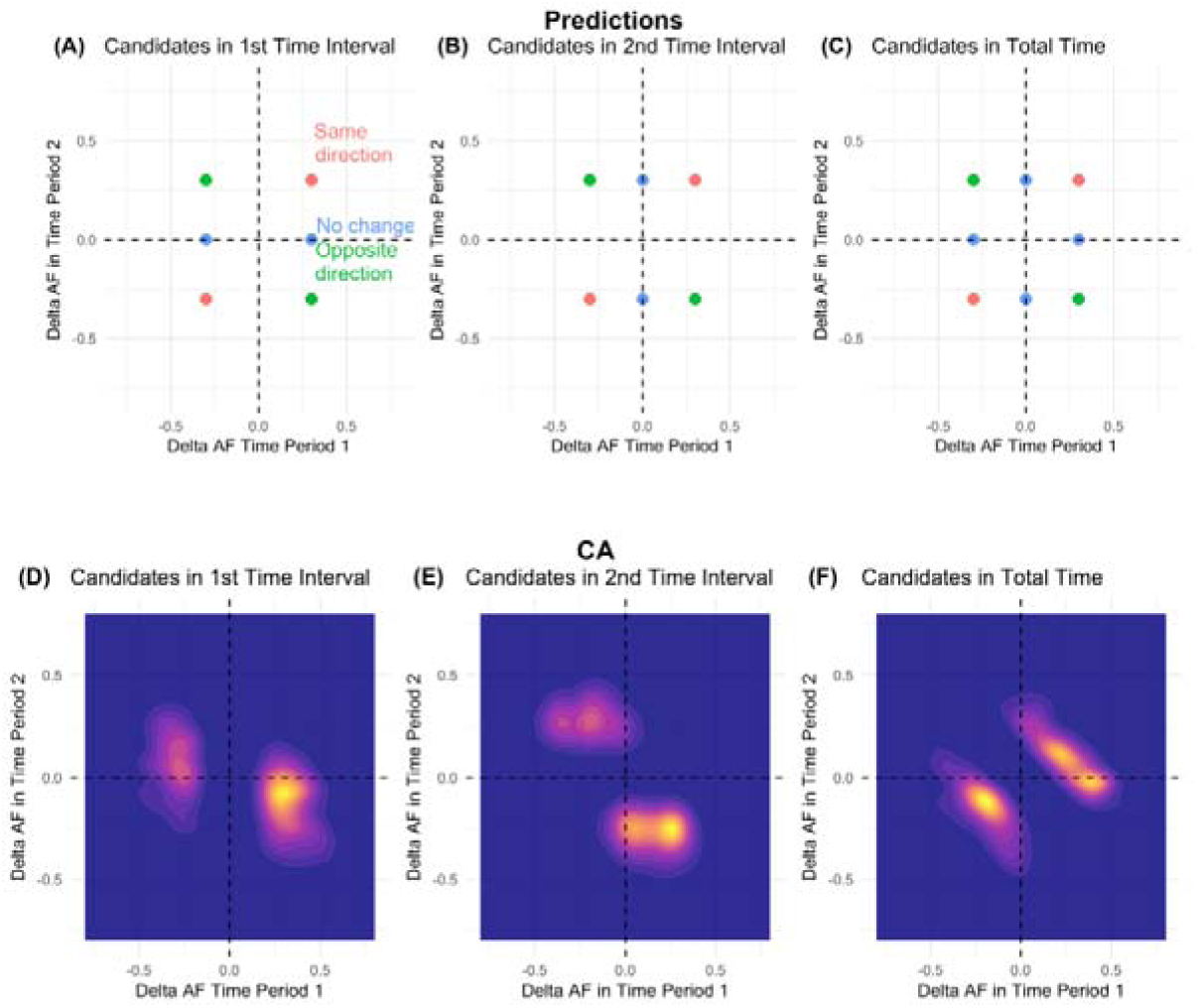
Allele frequency change dynamics of candidate windows between time interval 1 and time interval 2, Predictions are plotted. **(A-C)** coloured with the same pattern, if there was a change in one of the time intervals and not the other (blue), if there was a change in the same direction in both time intervals (red), and a change in opposite directions in both time intervals (green). Observed allele frequency density plot is shown for an example population CA (**D-F**). Where density of data is coloured from yellow (most dense) to purple (less dense). Other populations showed similar distributions and can be found in the Figure S12 – FIgure S14.

## Discussion

By using temporal genomic sampling we were able to uncover the dynamics of genome change in early adapting populations. We found more repeated signals of selection, and therefore more predictable genomic evolution, during the early stages of adaptation. Both chromosome– and window-level analyses revealed more shared outliers among our populations after roughly 10 generations than after roughly 20 generations. Also, genome-wide covariance between replicate population pairs was the highest in the first period. This pattern provides empirical support for theoretical predictions that selection should be strongest and therefore more predictable early in adaptation, when populations are furthest from their adaptive fitness peaks (Orr, 1998). This interpretation is further supported by documented, repeated, phenotypic evolution in these populations, where male colour pattern and size at maturity evolved similar differences between the translocated experiment and the source population roughly after generation 8.5 and 14 respectively (Kemp, Batistic and Reznick, 2018; Reznick *et al*., 2019). Similarly rapid phenotypic evolution has been reported in other translocation experiments (Reznick *et al*., 1997). This rapid and parallel genomic response may help explain why guppies serve as a powerful model system for studying convergent evolution.

Given our temporal genomics sampling design with replicate populations, we were able to directly measure selection response in both genome-wide patterns and at individual loci. There is some agreement in the dynamics between the two approaches; stronger repeatability across populations in the first period (as detailed above) and some evidence for fluctuating selection when comparing between time intervals. Specifically, candidate windows in the second period were more often in opposite directions from the first period, covariance between periods within a population was low genome-wide, and, for a few chromosomes, those covariances were significantly lower than expected under neutrality. Taken together, these results suggest that early adaptation involves both polygenic responses and responses at individual loci with large effects. This fits well with our association studies that show differences between natural LP and HP populations to be based on both polygenic and oligogenic sources (Whiting *et al*., 2021, 2022).

To our knowledge, this is the first study to explicitly test for a polygenic selection response in guppies following a change in predation regime. When compared to lab-based evolve-and-resequence studies, which often include more than ten replicates and organisms with short generation times, allowing for many more temporal sampling points, our power to detect a response is limited, (e.g. (Barghi *et al*., 2019)). However, when compared to other natural or semi-natural studies we have a similar number of temporal sampling points and number of replicated populations ((Saleh *et al*., 2022; Reid, Star and Pinsky, 2023; Tisthammer *et al*., 2025), but see (Lynch *et al*., 2024; Han *et al*., 2025) for more temporal sampling points). Most of these studies report a positive genome-wide covariance over time within populations and therefore conclude that the response to selection was polygenic. We also report positive covariances within populations over time but crucially these are within the range of covariances simulated under neutrality. We have the unique advantage of using known census size to inform our simulation models from mark-recapture studies, however, simulations using estimated parameters are also informative. For example, Han et al. (Han *et al*., 2025), similarly report a positive covariance over time in cod but reject the hypothesis of a selection response as it falls within neutral simulations. We expect that the positive covariance values under neutrality are due to either the relatively short time period or strong linkage disequilibrium with selected SNPs. Overall, we caution against interpreting covariance patterns without the support of simulations.

Given the number of population replicates we had the ability to test for genome-wide covariance within a period between populations and therefore the relative repeatability of polygenic response over time. It is striking that in the first time interval, between-replicate covariances exceeded the 90th percentile of simulated covariance under neutrality, and even without chromosome 15 that contains a large-effect locus, covariance estimates were above the 80th quantile. The highest values were observed in replicate pairs sharing similar starting genetic diversity (CA–TA and LL–UL), as well as in one pair exposed to a common light environment (TA–UL) (Reznick *et al*., 2019), as would be predicted under a common genomic selection response. The covariances were smaller in the second time interval but still higher than the mean of simulated covariances, within the 80th quantile. Shared source population can explain positive covariance among populations under neutrality but a parallel selection response is responsible for the higher than simulated covariances found here.

Through chromosome-wide covariance between population and overlapping window analysis we uncovered targets of selection. We found a number of chromosomes with covariances between populations that exceeded the magnitudes estimated from simulated distributions. This is strong evidence for linked parallel selection. Chromosome 15 is the most consistent outlier, likely driven by a region at 5Mb, a candidate window found here and previously documented as showing signals of selection (Van Der Zee *et al*., 2022). In addition, we detect new signals for repeated selection on chromosomes 1 and 21 in the first period and chromosome 20 in the second. Notably, a large haplotype (11-14Mb) on chromosome 1 has been linked to colour variation in other populations of guppies (Paris *et al*., 2022) and a region on chromosome 20 at 2Mb has been linked to repeated divergence between natural high and low-predation population pairs (Whiting *et al*., 2021). However, neither of these specific windows show significant allele frequency change in our window-based analysis. A possible explanation for this discrepancy is that selection is acting on different haplotypes but with the same beneficial alleles, e.g. soft selective sweeps (Barrett and Schluter, 2008). Soft selective sweeps can produce subtle yet parallel shifts in allele frequencies that are difficult to detect using standard window-based approaches. Selection signals found at known genes involved in colour and appetite also provide good candidates for further inquiry.

Our unique, ongoing monitoring of these populations also allowed us to examine the effect of demographic events on genomic diversity. We found signals of bottlenecks likely caused by founding events are no longer apparent approximately 20 generations after transplantation, despite documented declines in census population size. Across both periods, we detected no significant reduction in nucleotide diversity. However, after the first period, all populations exhibited longer and more frequent runs of homozygosity (ROH) compared to the source population. This pattern is consistent with strong inbreeding during early adaptation, rather than an overall loss of genetic diversity. In contrast, although most populations experienced a severe crash during the second period due to a flooding event (month 106), size and number of ROH decreased across all populations and became comparable to levels observed in the source. The population crash was followed by rapid population recovery, which likely explains the absence of a lasting genomic signature. We also detected persistent genome-wide differentiation among our populations, likely driven by differences in fluctuations early in their history. Specifically, pairwise F_ST_ values involving TA are the highest at both time points (2013 and 2018) and this may be attributable to an earlier and longer population crash when numbers were already low (month 31 – 37) that was not observed in other populations. Overall, our results show that population declines occurring at different points in a population’s history are likely to have different genomic consequences. This distinction will also be useful for those exploring the genomic effects of extreme climate events, such as the large flooding observed here, in the conservation of vulnerable species (Heinen *et al*., 2025).

In summary, we document rapid and predictable evolution in response to a novel environment in as little as 10 generations. The majority of the repeated change occurs early, highlighting the importance of initial adaptive responses. While at the phenotypic level, rapid evolution is now accepted, with the guppy system providing some of the earliest and most compelling evidence, this study shows an early genomic response, a process only beginning to be appreciated by population geneticists (Messer, Ellner and Hairston, 2016). Crucially, using multiple time points we were able to document the dynamics of both neutral evolution and selection.

## Acknowledgements

We would like to thank Josie Paris, and Reid Brennan for discussion and thoughtful comments on the manuscript. We would like to acknowledge the high performing computing (HPC) ISCA server at the University of Exeter. We would also like to thank the Ramlal family in Trinidad for providing housing and logistical support for the field research as well as the hundreds of Young Research Scientists involved in collection of field data. This work was supported by an EU Research Council grant (GuppyCon 758382) BAF, the National Science Foundation USA (DEB-0623632EF, DEB-0808039, DEB-1258231, DEB-1556884, DEB-2247042) to DNR, JT, and RB.

## Data and Code Accessibility Statement

The data that support these findings are openly available at: European Nucleotide Archive (https://www.ebi.ac.uk/ena/browser/home)—reference numbers: PRJEB42705 (all experimental populations sampled in 2013) and PRJEB10680 (GHP) reads for data sampled in 2018 found here (PRJEB106412). VCF file for final data can be found on figshare (10.6084/m9.figshare.32604111). Code for SNP calling and read processing found here (https://github.com/josieparis/gatk-snp-calling). Code for all other analyses can be found here (https://github.com/bfraser-commits/Temporal-genomics).

