## Supplementary figures for "Temporal genomics reveals rapid parallel adaptation in experimental populations of Trinidadian guppies (*Poecilia reticulata*)"

**Figure S1:** Correlation between depth and covariance and variance per chromosome

**Figure S2:** PCA of source and experimental populations for PC3 and PC4.

**Figure S3:** Tajima’s D genome-wide violin distributions for the source population (GH) and each introduction at both time periods.

**Figure S4:** Expected heterozygosity genome-wide violin distributions.

**Figure S5:** Frequency of ROHs per size class per population

**Figure S6 - Figure S8:** Allele frequency covariances per chromosome among populations for each time interval and within population across the two time interval

**Figure S9:** Chromosome size and covariance correlations examples

**Figure S10:** Genome-wide allele frequency covariance without chromosome 15 for each time interval and within population

**Figure S11:** Comparison of simulated and observed allele frequency change modelled within windows across time periods, dashed lines indicate 95% cut-off used on observed data to identify selected sweeps.

**Figure S12:** Allele frequency change dynamics of selective sweeps between time period 1 and time period 2 for population TA

**Figure S13:** Allele frequency change dynamics of selective sweeps between time period 1 and time period 2 for population UL

**Figure S14:** Allele frequency change dynamics of selective sweeps between time period 1 and time period 2 for population LL

**Table S1**: Sampling information and mean sequencing coverage

**Table S2**: Genome-wide mean and SEM of FST of introductions by year and GH estimated in 50kb windows.

**Table S3**: Pairwise genome-wide FST

**Table S4**: Genome-wide diversity statistics (pi, Tajima's D, and Heterozygosity) and comparisons with the source population

**Table S5**:Table S5: Runs of homozygosity in each population, and FROH comparison with the source population

**Table S6**: Outlier window analysis for each population

**Table S7**: Candidate windows per time interval

**Table S8**: Gene annotation for candidate windows in the first time interval

**Table S9**: Gene Annotation for candidate windows for the second time interval

**Table S10:** Gene Annotation for candidate windows for the total time interval

**Table S11**: GO enrichment for candidate windows in Time period 1, Time period 2, And Total time period

**Table S12:** Simulated cut-offs of allele frequency cut-offs forward simulating a VCF

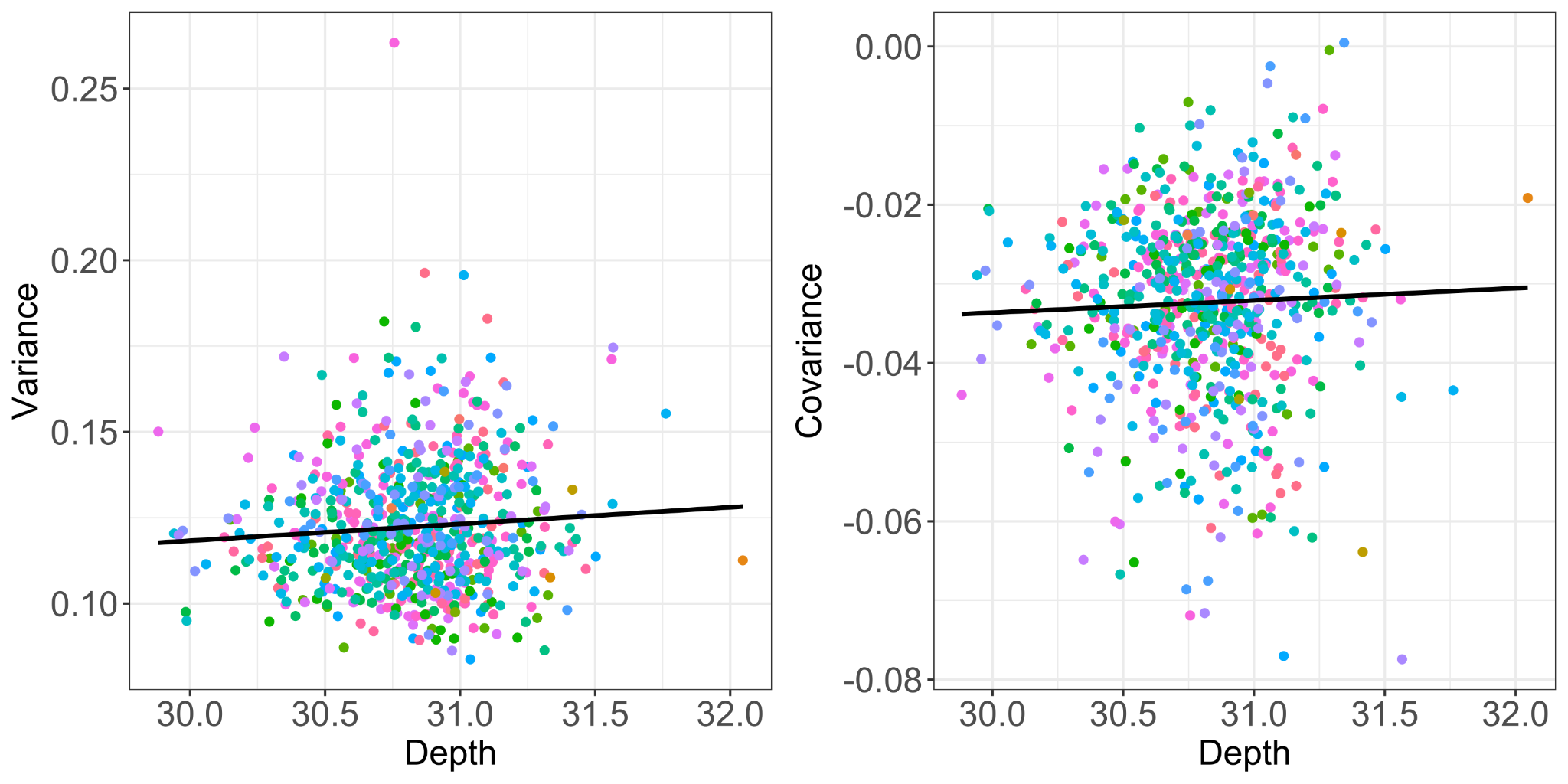

**Figure S1**: Diagnosis plots for cvtk. Showing depth of coverage against variance and covariance. Colours indicate chromosomes.

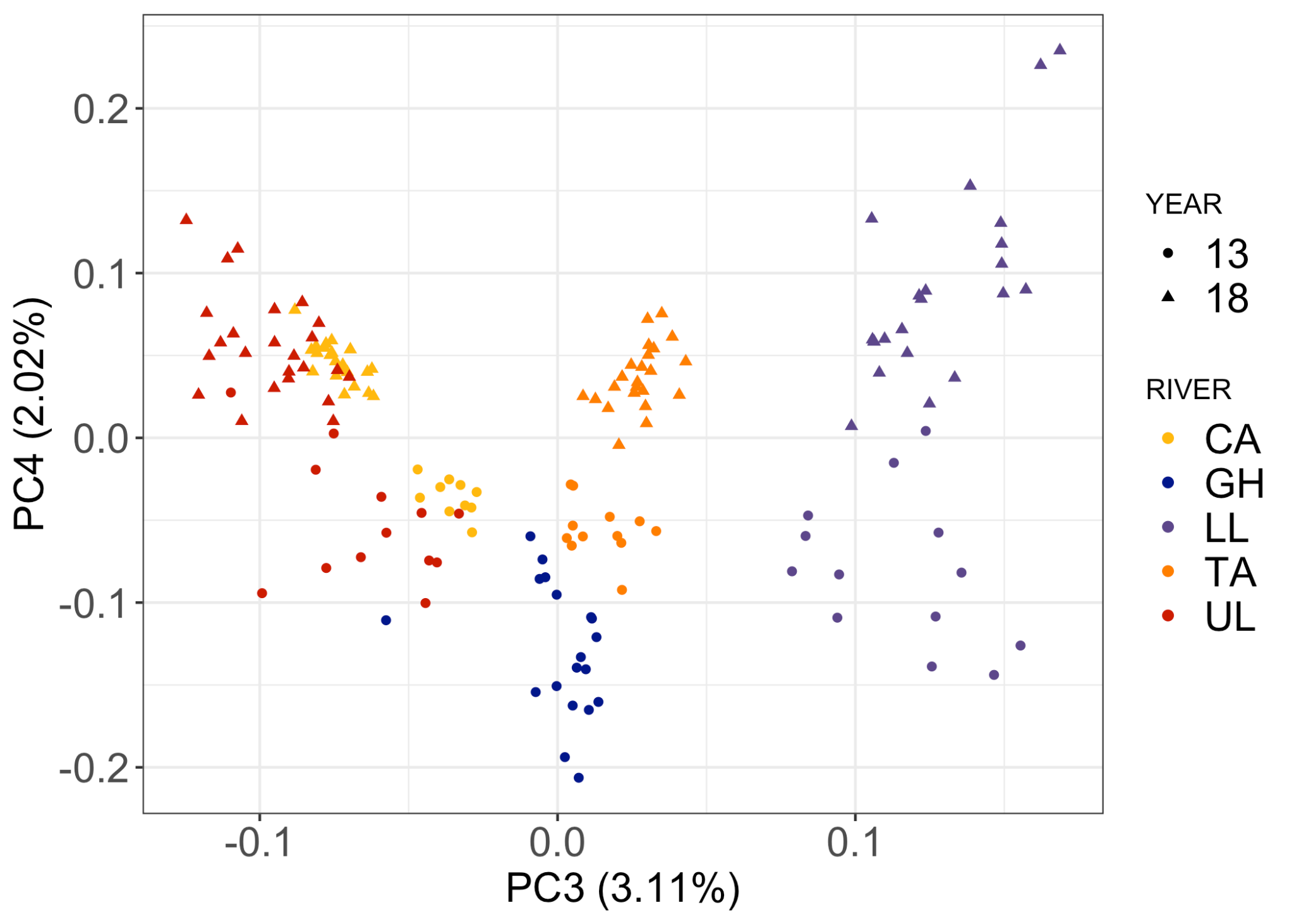

**Figure S2**: PCA for populations and time periods, showing PC3 against PC4.

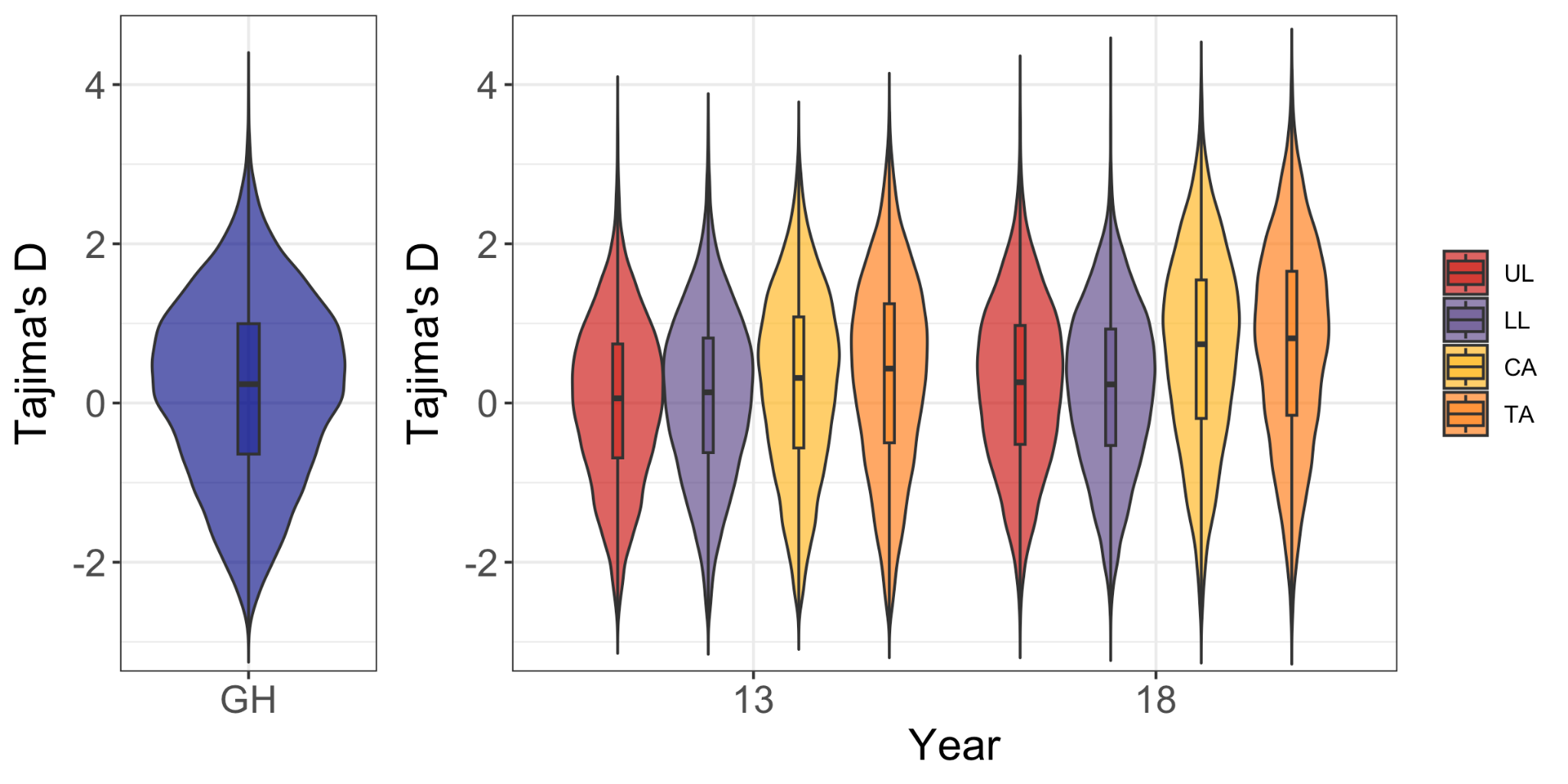

**Figure S3:** Tajima’s D genome-wide estimates violin plots per population per time period.

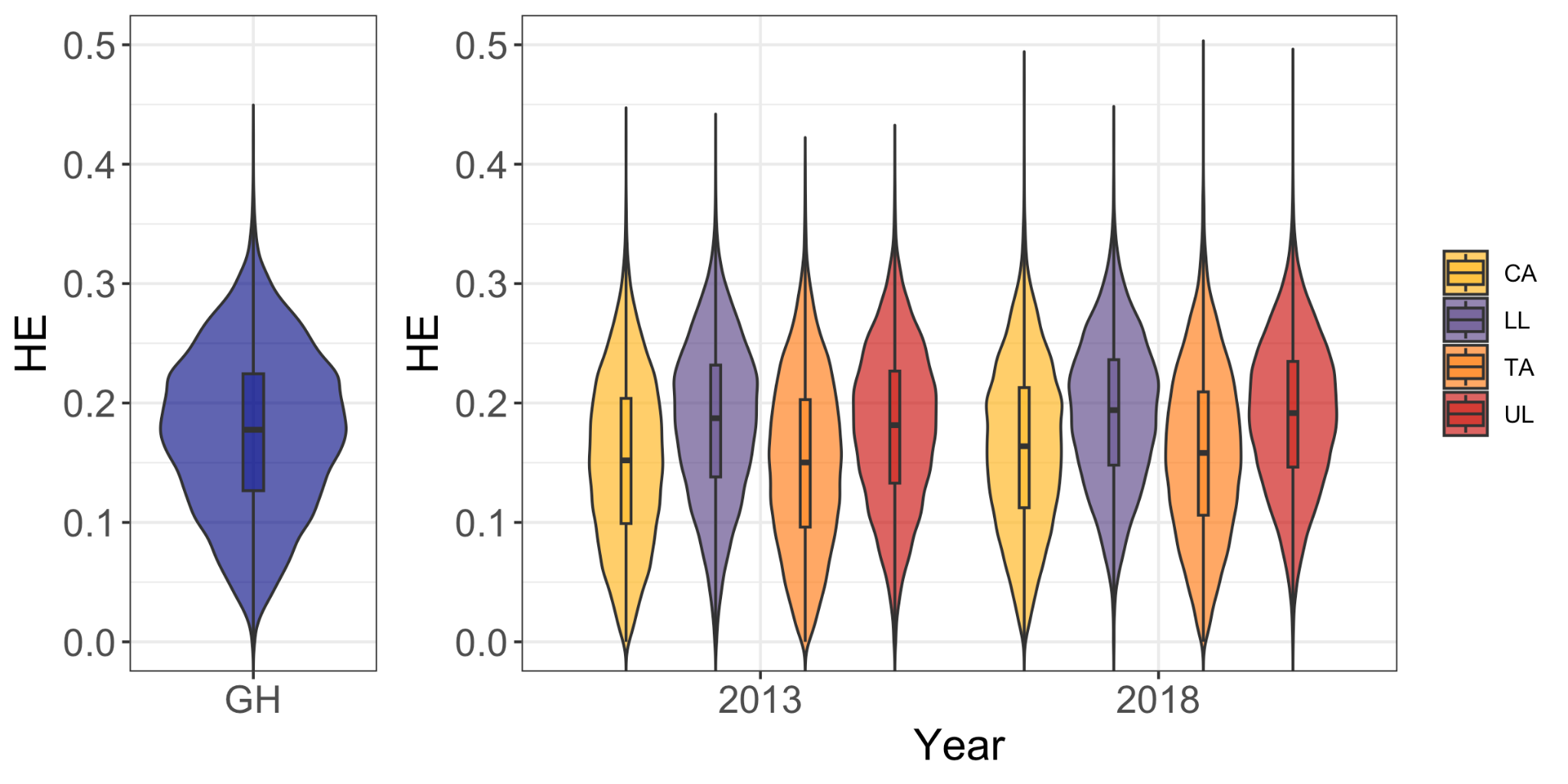

**Figure S4:** Expected heterozygosity genome-wide estimates.

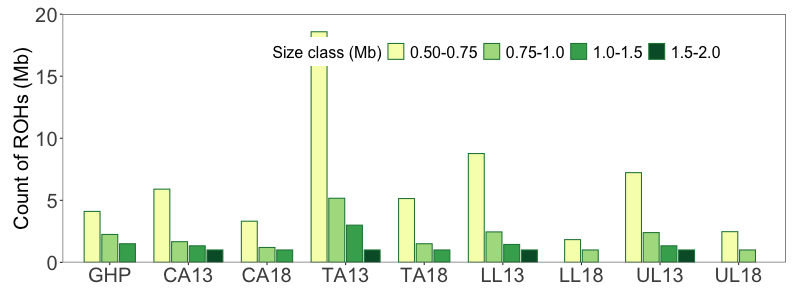

**Figure S5:** Count of ROHs per population per size class.

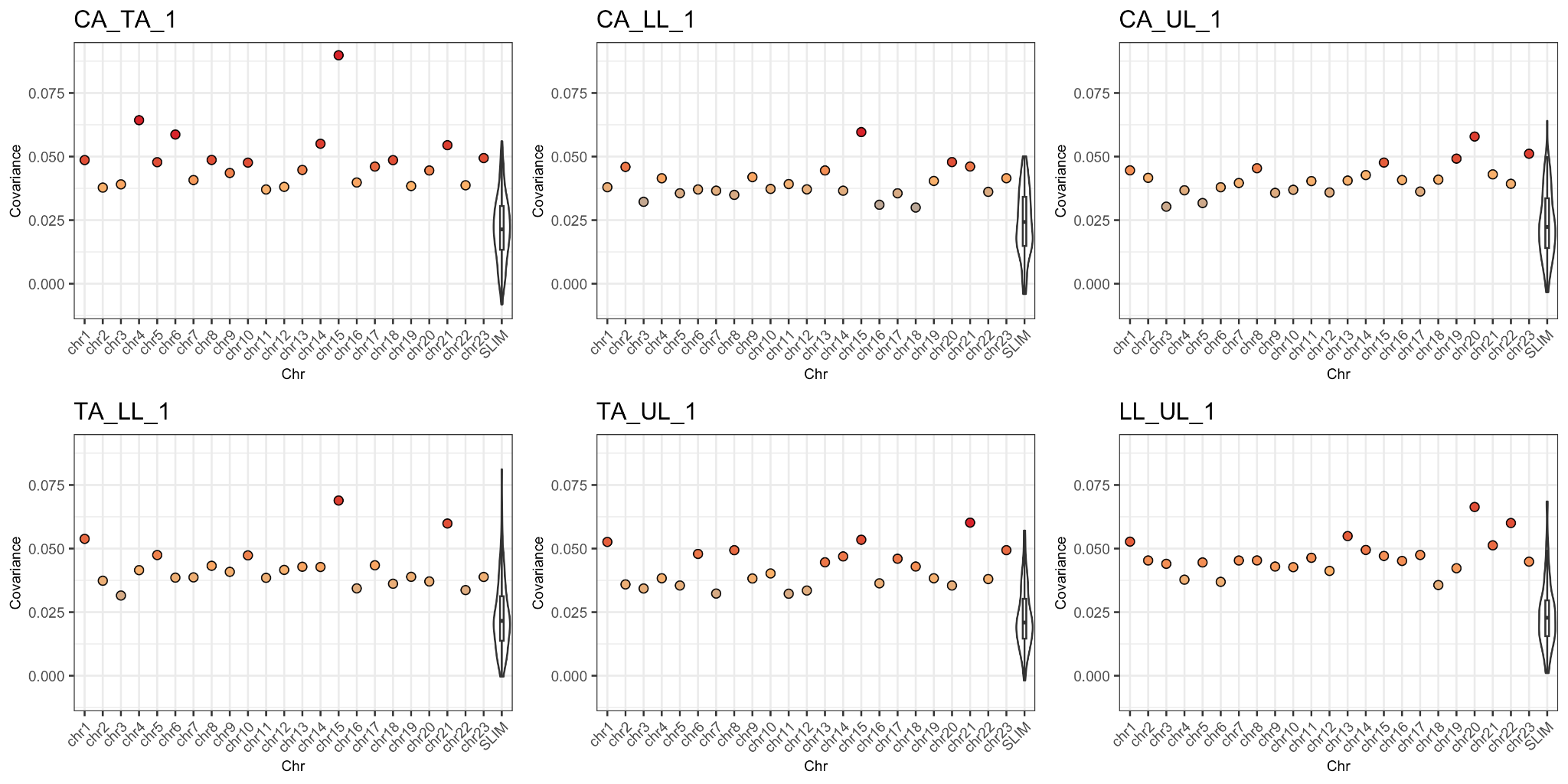

**Figure S6**: Genome wide covariance between pairs of populations within the first time interval, per chromosome (coloured points) compared to simulated distributions (violin plots).

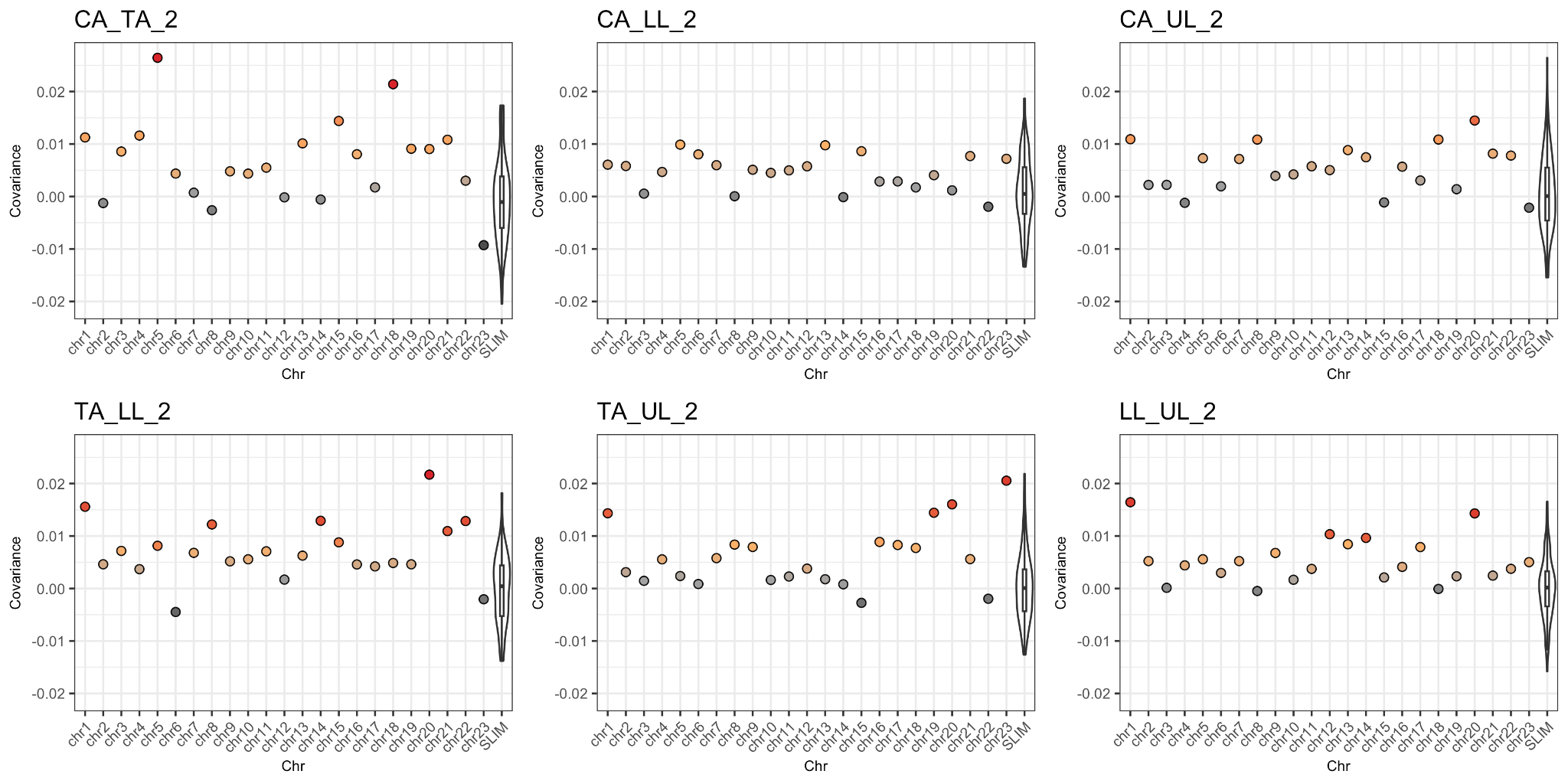

**Figure S7:** Genome wide covariance between pairs of populations within the second time interval, per chromosome (coloured points) compared to simulated distributions (violin plots)

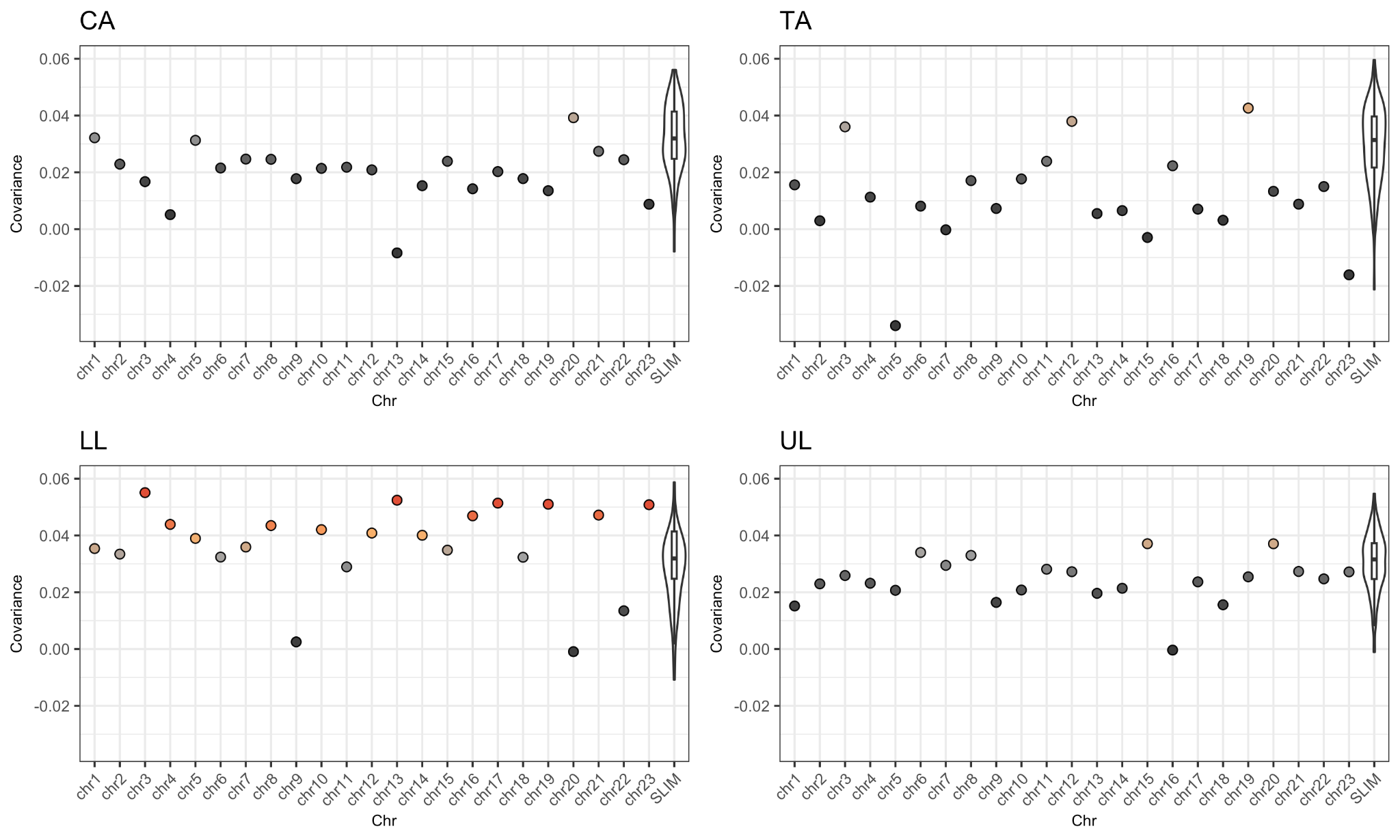

**Figure S8:** Covariance between the two time intervals for each population, per chromosome (coloured points) as compared to simulated distribution (violin plots)

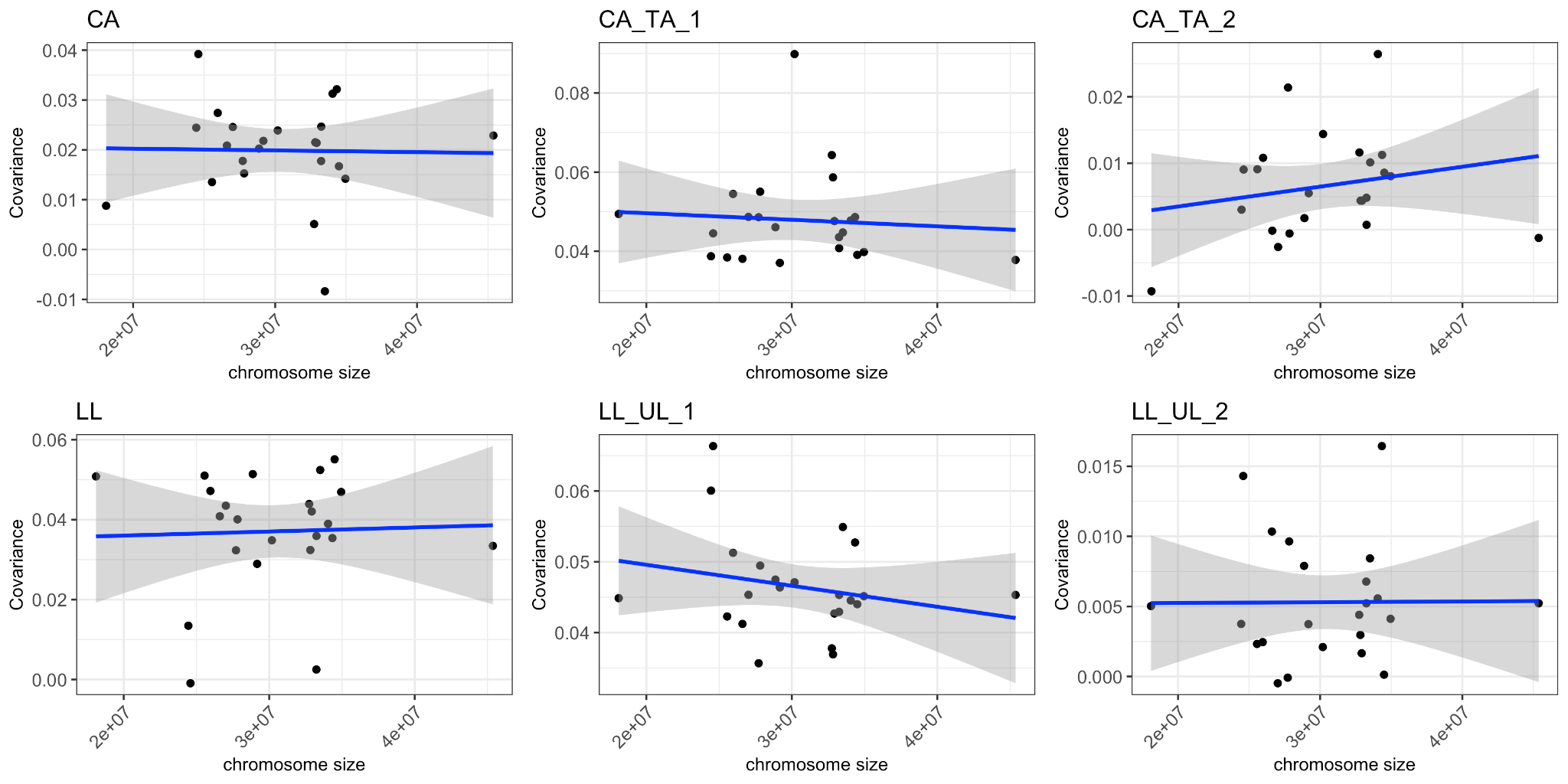

**Figure S9:** Example of chromosome and covariance correlations for 6 of the 18 comparisons.

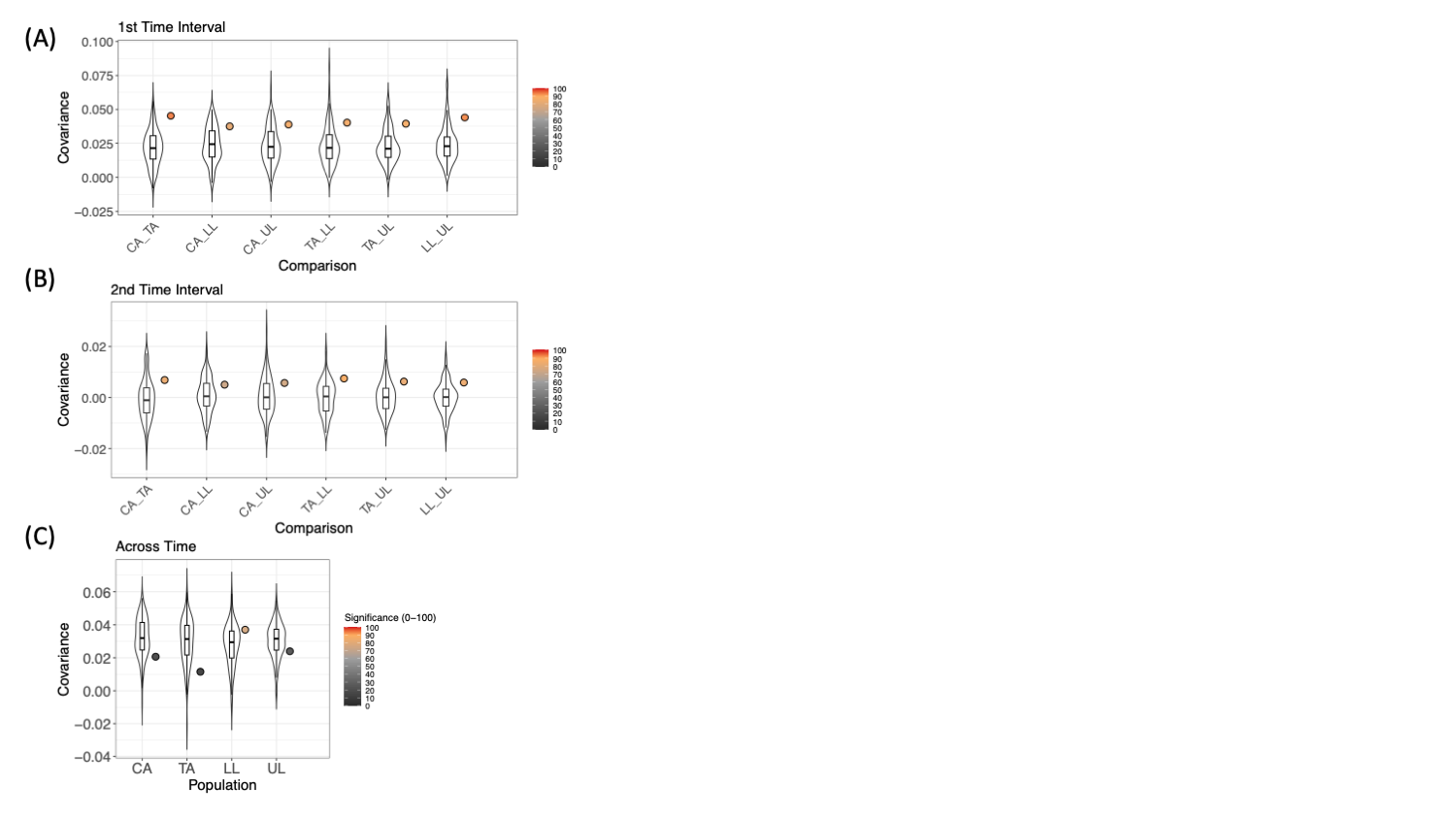

**Figure S10:** Genome-wide covariances without chromosome 15, observed in coloured points and simulated distributions in violin plots.

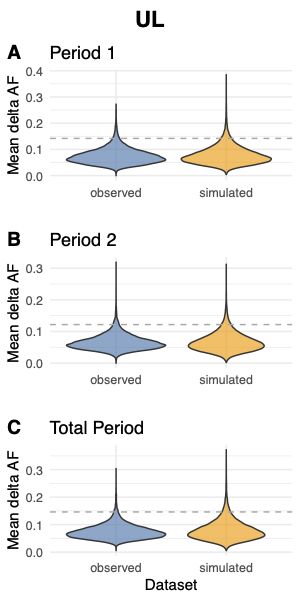

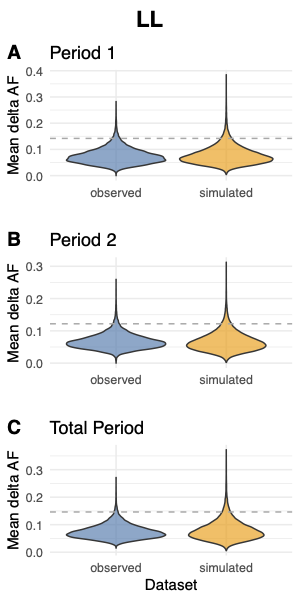

**Figure S11:** Comparison of simulated and observed allele frequency change modelled within windows across time periods, dashed lines indicate 95% cut-off.

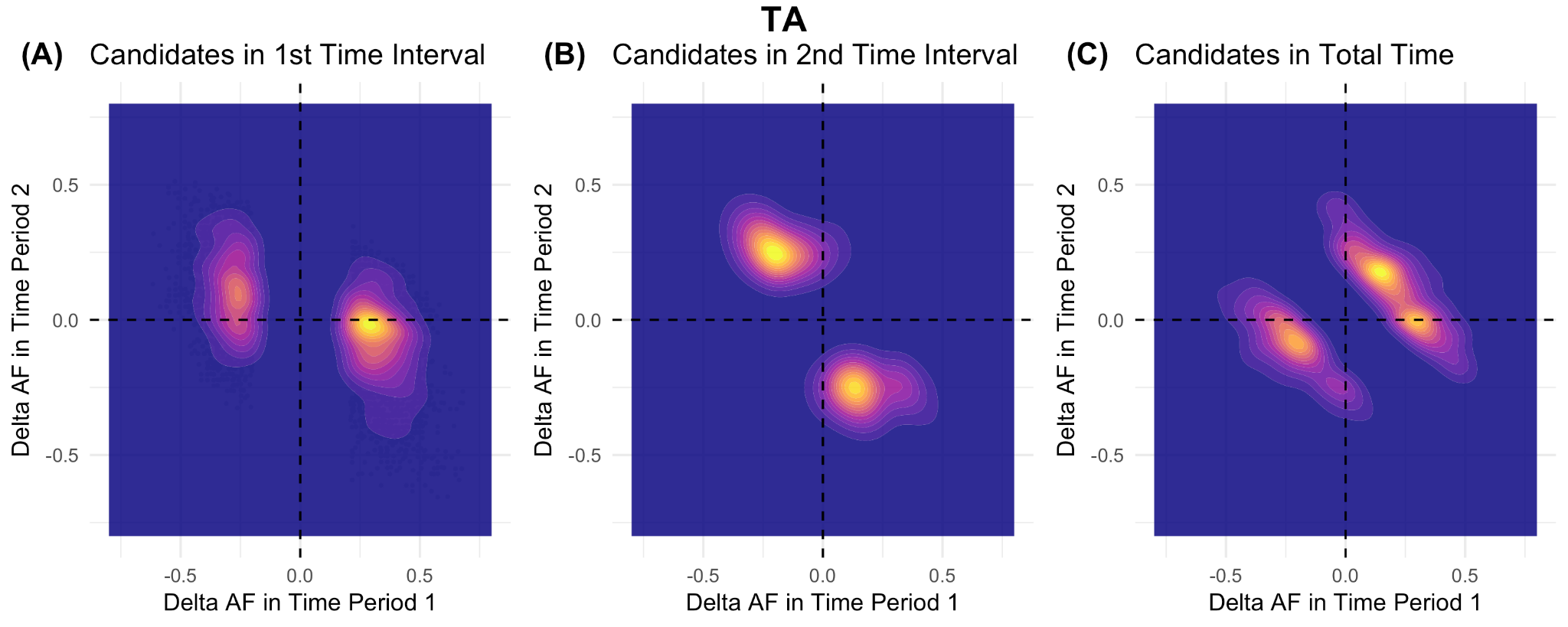

**Figure S12: Allele frequency change dynamics of selective sweeps between time period 1 and time period 2 for TA.** The density of data is coloured from yellow (most dense) to purple (less dense).

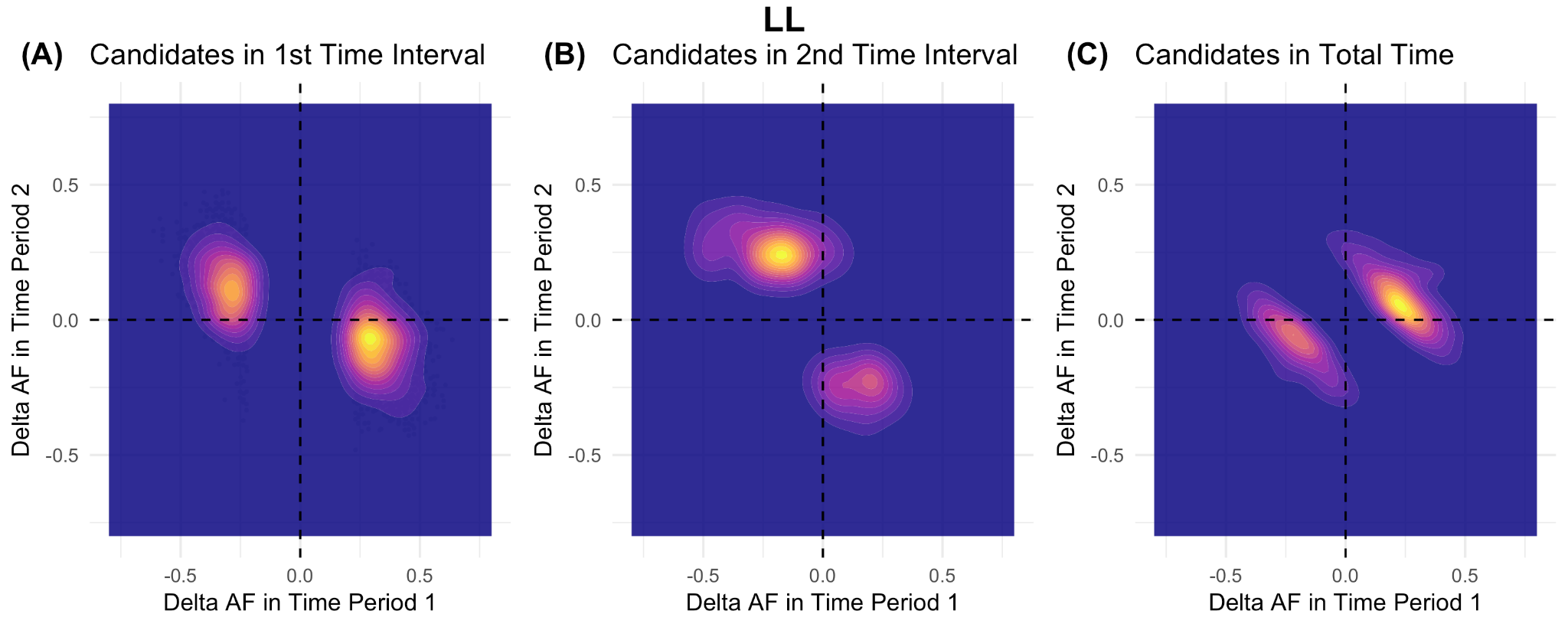

**Figure S13: Allele frequency change dynamics of selective sweeps between time period 1 and time period 2 for LL.** The density of data is coloured from yellow (most dense) to purple (less dense).

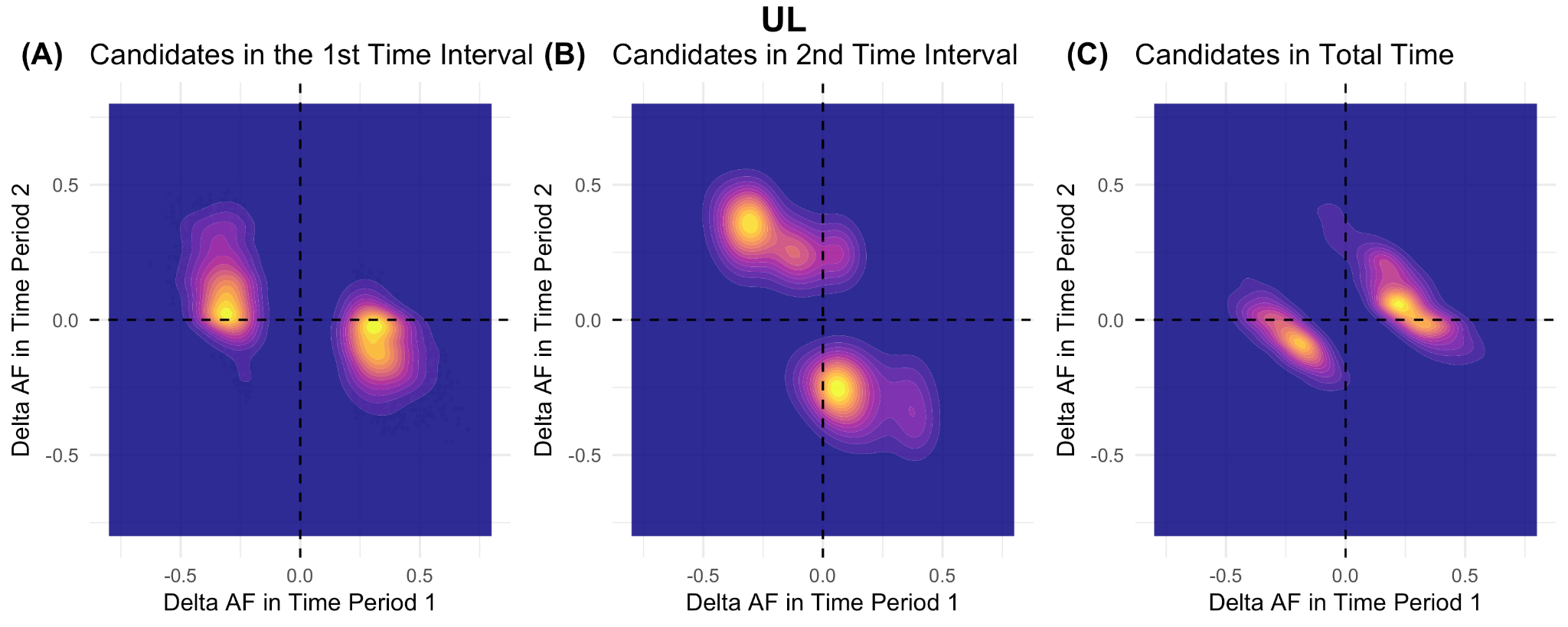

**Figure S14: Allele frequency change dynamics of selective sweeps between time period 1 and time period 2 for UL.** The density of data is coloured from yellow (most dense) to purple (less dense).
